# Direct mechanical communication of cellular to nuclear shape in oocytes

**DOI:** 10.1101/2025.02.17.638579

**Authors:** Bart E. Vos, Yamini Vadapalli, Till Muenker, Ida Marie Astad Jentoft, Elena Todisco, Mohammad Amin Eskandari, Melina Schuh, Peter Lenart, Timo Betz

## Abstract

The mechanical properties of the cytoplasm and nucleoplasm are crucial for the correct and robust functioning of a cell and play a key role in understanding how mechanical signals are transferred to the nucleus. Here, we demonstrate remarkable shape mimicry between the cellular and nuclear shape of oocytes, following the externally applied deformation without direct contact between the cell cortex and the nucleus. This effect arises from a surprisingly soft and fluid-like nucleoplasm that is barely resisting any external strain, while the viscoelastic cytoplasm drives shape transmission. Comparative studies in jellyfish, starfish, and mouse oocytes reveal that lower cytoplasmic elasticity in jellyfish leads to reduced nuclear shape mimicry, highlighting the role of cytoplasmic mechanics in nuclear deformation.

**Significance Statement:** Mechanosensing of the nucleus is, in addition to chemical signalling, an important factor in gene expression. Although nuclei are often thought to be rigid inclusions in the cytoplasm of a cell, we show that in oocytes nuclei are much more deformable. Using a combination of intranuclear, intracellular and extracellular measurements, we attribute our findings to a fine balance between the soft nucleoplasm surrounded by an elastic shell and the viscoelastic properties of the cytoplasm.

## Introduction

The nucleus of a cell is arguably its most important organelle, but definitely its largest. Hence it not only plays an essential role in the storage of genetic material, it is also an important contributor to the mechanical properties of the cell as a whole. In turn, mechanical deformations of the nucleus (as well as other organelles^1^) play an important signaling role, which has been an emerging research topic in the past decades.^2–5^

To date, nuclear mechanical properties have been investigated using a wide range of techniques, ranging from whole cell deformations,^6–10^ and local force applications on the single cell level,^11–13^ to direct interaction with the nucleus via osmotic nuclear compression,^14, 15^ micropipette aspiration of isolated nuclei,^12, 16, 17^ atomic force microscopy^18^ or tracking of intranuclear particles.^19^ Despite this wide variety of techniques used on a range of different cells, all studies find that the nucleus is significantly stiffer and less deformable than the remainder of the cell.

However, only few studies have measured the mechanical properties inside the nucleus, mainly due to technical difficulties. Magnetic tweezers have been used to investigate intranuclear mechanics.^20–22^ While this technique has limited temporal resolution and the precises forces acting on the probe particle are difficult to determine, it provides the first active rheology insights into the nucleus. More recently, optically tweezers have been used to insert probe particles in the nucleus and perform active intranuclear microrheology.^23^ However, as the nucleus is a multi-component organelle with distinct properties,^3, 4, 24^ a full mechanical characterisation requires knowledge of the intranuclear properties over a wide range of biologically relevant timescales.

Non-invasive techniques have also been used to measure intranuclear mechanics.^19, 24–26^ However, the nucleus cannot be considered a purely passive environment,^22, 27, 28^ and the combination of Brownian and active motion leads to more fluctuations and thus an underestimation of the mechanical properties using passive interpretation approaches alone.^29, 30^ A novel approach based on Onsager’s regression hypothesis may overcome this.^31, 32^

In this study we investigate the intracellular and intranuclear mechanics of oocytes. While most studies regarding nuclear mechanics have addressed somatic cells, one of the most fundamental cell types, the oocyte, have been hardly investigated. Oocytes are exceptionally large cells, which store nutrients to support embryo development after fertilization. As they also store nuclear proteins, the nucleoplasm also has an exceptionally large volume. This results in a specialized nuclear architecture, with a large amount of nucleoplasm and relatively small chromosomes scattered in this large volume. In somatic cells the nucleus is filled mostly with chromosomes (chromatin) and there is little nucle-oplasm. Therefore, oocytes provide a unique opportunity to investigate the properties of the nucleoplasm, separately from chromatin.

Since oocytes are the origin of any developing organism, any permanent damage to the nucleus can lead to catastrophic defects, which suggests that the nucleus should receive special protection against external force. Furthermore, during maturation and gonad expulsion, oocytes are subject to large external forces, again suggesting the requirement of efficient mechanical protection for the nuclei of these fundamental cell types. Recently, the first detailed framework for ovulation has been reported, confirming a significant deformation during expulsion from the follicle.^33^

To test such potential shielding mechanisms, we quantify the deformability in response to an external compressive force of starfish, jellyfish and mouse oocytes and their nuclei, covering a good part of animal phylogeny. To understand the surprising finding that the nucleus closely mimics the overall cell deformation, we determine the mechanical properties of the nucleus and the cytoplasm by active, optical tweezers based microrheology on micron-sized probe particles. In contrast to studies on other cell types we find that the oocytes’ nucleoplasm is more than an order of magnitude softer than the cytoplasm. Our measurements suggest that in oocytes the nucleoplasm is not shielded from external deformations by a hard-to-deform nucleoplasm or by a lamin network that lines the interior of the nuclear envelope, but rather follows whole-cell deformations where the forces are transmitted by the soft, viscoelastic cytoplasm. Although this sounds counter-intuitive given the importance and potential sensitivity of the oocyte nucleus, this open the door for new mechanosensing events, as any external force acting on the oocyte is rapidly reflected in the nuclear shape.

## Results

### Nuclear shape follows cell shape during cellular compression in starfish oocytes

An oocyte is exposed to variable forces after its expulsion from the gonad. In order to investigate if the nucleus is mechanically shielded from external forces, or if it is directly sensing such forces, we decided in a first experiment to mechanically challenge a starfish oocyte and register the effect on the nucleus. In detail, the potential nuclear deformation is measured after applying a compressive force on a jellyfish oocyte by pushing the oocyte against the side of a glass coverslip with a blunt indentor (Fig. 1a). To our surprise, at first sight the cell fails to shield the nucleus from the extracellular deformation, as the nucleus adopted a shape similar to the shape of the deformed cell (Fig. 1b,d).

**Figure 1:**
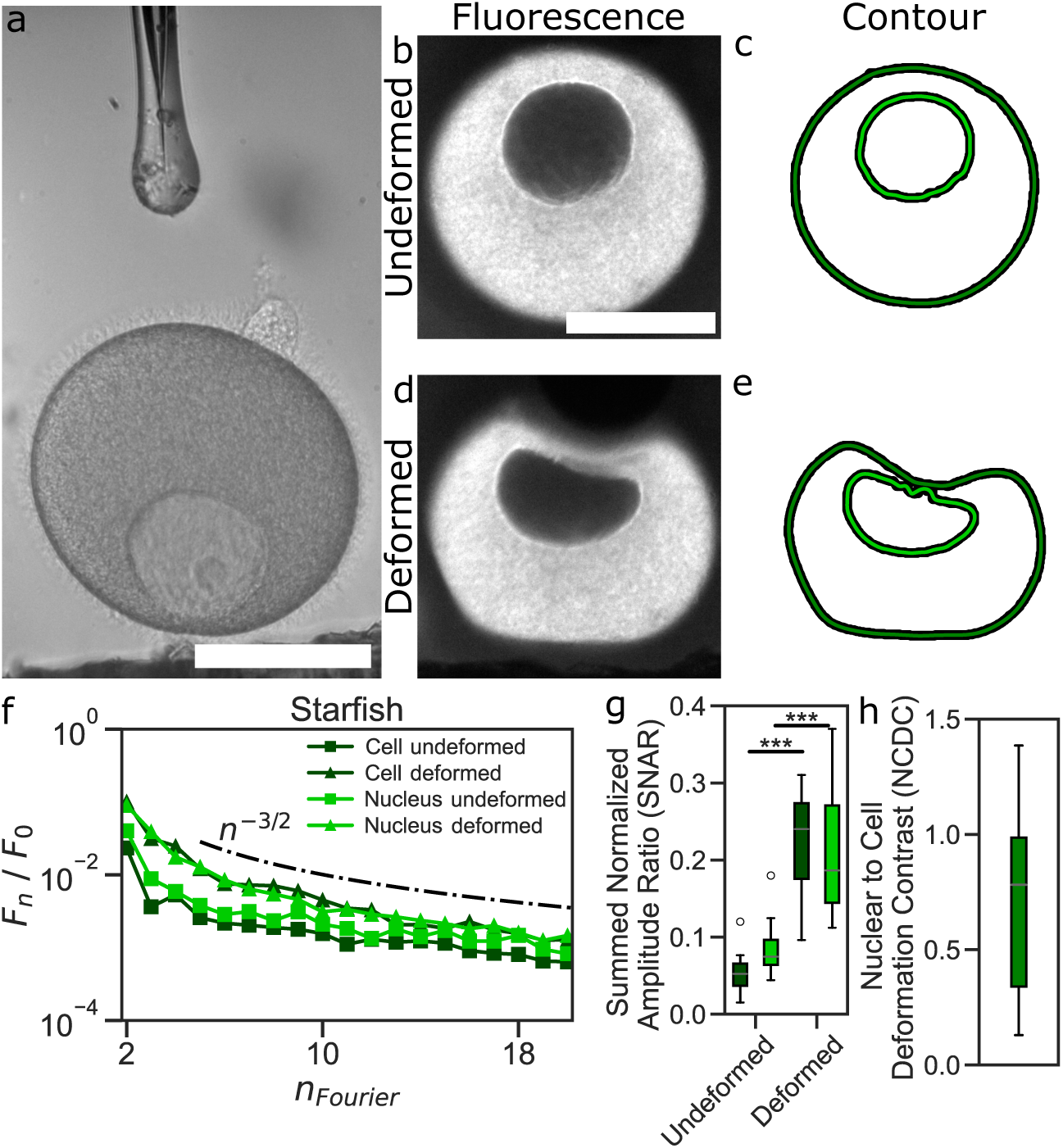
Analysis of cell and nuclear deformation. **a**: Brightfield image taken prior to deformation of a starfish oocyte, **b,d**: fluorescence images before and during deformation, **c,e**: contours of the oocyte and the nucleus before and during deformation (black dots), overlaid with a fit using the zeroth to the nineteenth Fourier modes (solid green lines) of the contour, **f**: average Fourier mode 2 to 19 for the cell and nucleus of a starfish oocyte, with power law decays (dashdotted line: *n*^−3*/*2^), **g** the Summed Normalized Amplitude Ratio (*SNAR*) for starfish oocytes, and **h** the Nuclear to Cell Deformation Contrast (*NCDC*) for starfish oocytes. Scale bars: 100*µ*m. Black lines represent the standard deviation. ***: *p <* 0.001.

To better determine the shape change that this compression induces in the cell and the nucleus, we decided to develop a quantitative way to compare the shapes of the cell and the nucleus. To this end we first obtained the contours of the cytoplasm and the nucleus from microscopy images (Fig. 1c,e), which confirm the initial impression of similar shapes. For a more quantitative analysis of these deformations we exploit the length scale decomposition of the contour into Fourier modes. This was obtained by defining the pixel-based center of mass of each contour, allowing extraction of the angle-dependent contour radius and finally calculation of its Fourier coefficients. The power of this approach is that a length-dependent separation of any simply connected shape is reached. Originally introduced for vesicles ^34^ the Fourier decomposition is independent of the orientation, and insensitive to lateral displacements or errors in the center of mass inference as these only influence the first mode. Other quantification approaches focus on nuclear elongation only,^35^ but might not appreciate the complexity of the deformed cell and nuclei. The aspect ratio or a correlation analysis works well for low deformations but either breaks down at high deformations or is sensitive to the orientation and does not yield information on the similarity with respect to different length scales.

In the Fourier decomposition, a perfect circle only has a zeroth order Fourier mode *F*_0_ that reflects its radius, and the first mode is only different from 0 when the center was not chosen optimally. In consequence, the differences to a perfect circle are encoded in the higher order modes. The effect of the contours’ global sizes can be conveniently removed by normalizing all modes by the *F*_0_ mode. This is reflected in Fig. 1f, showing up to Fourier mode twenty for an undeformed and a deformed starfish oocyte.

As already seen in the visual inspection of Fig. 1, we find a quantitative resemblance of the modes between the undeformed and deformed cell and nucleus. Additionally, we find that the values of the higher modes decay rapidly, suggesting that the relevant information is already captured by the lower modes. Interestingly, the decay of the Fourier amplitudes with mode number seems to be consistent with a power law of exponent −3/2 for higher modes, that is close to values reported in active vesicles.^36^

To characterize the cellular and nuclear deformation by a simpler quantity that reflects the strength of the Fourier modes we define the Summed Normalized Amplitude Ratio (*SNAR*) as the sum of mode 2 to 19 normalized by mode 0 to obtain a size-independent measure:

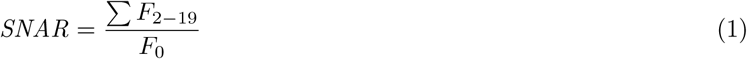

The upper limit of 19 was chosen as this is approximately equal to the resolution of the microscope. With vanishing *SNAR ≈* 0, the cell or nucleus is perfectly round, while *SNAR >* 0 corresponds to some degree of deformation. We find that the *SNAR* remains close to 0 for the contour of the undeformed starfish oocytes (*SNAR^cell^* = 0.067), confirming that their unperturbed shape is a near-perfect circle (Fig. 1g).

While the *SNAR* gives an excellent quantification of a deformed shape, it does not provide a direct comparison between the nuclear and cellular deformation. To better address this resemblance we introduce the Nuclear to Cell Deformation Contrast (*NCDC*) which can vary between 0 (no nuclear deformation upon cell deformation) to 1 (same nuclear and cell deformation upon cell deformation), while values larger 1 mean that the nucleus deforms more than the cell. The *NCDC* is defined by the difference between the deformed and undeformed *SNAR* of the nucleus divided by the same quantity of the cell:

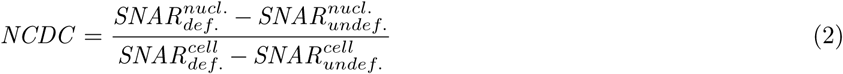

The *NCDC* quantifies how susceptible the nucleus is to deformation with respect to the cell, independent of the size of the cell and the nucleus and the initial and total degree of deformation. Indeed the *NCDC* of starfish oocytes shows with a value of 0.6 that the nuclear deformation partially recapitulates the shape of the cell upon deformation (Fig. 1h). This unexpected behaviour suggests that such resemblance in shapes might be an important feature of oocytes.

### Nuclear shape adaptation across different species

To test whether the here-observed shape resemblance is particular to starfish oocytes or can also be found in other species, we turned to jellyfish, a species that shares the overall environment with the starfish, but comes from to a very different evolutionary origin. If there is a systematic advantage for such a shape mimicry in free floating oocytes in sea water, we would expect a similar behaviour.

However, following the same experiments, we find that the *NCDC* of jellyfish is much closer to 0, indicating that the nucleus does not follow the shape of the overall cell, as can be seen in the *NCDC* values (Fig. 2a). To understand this better, we inspected possible molecular differences between the nuclei of starfish and jellyfish oocytes that might be related to their shape. A prominent candidate for such biomechanical differences are actin filaments. Indeed, while the nucleoplasm of starfish oocytes does not contain a concentration of actin comparable to the cytoplasm, this situation is reversed in jellyfish. Here the nuclear f-actin as stained by Utrophin is even larger than in the cytoplasm (Fig. 2b-c).

**Figure 2:**
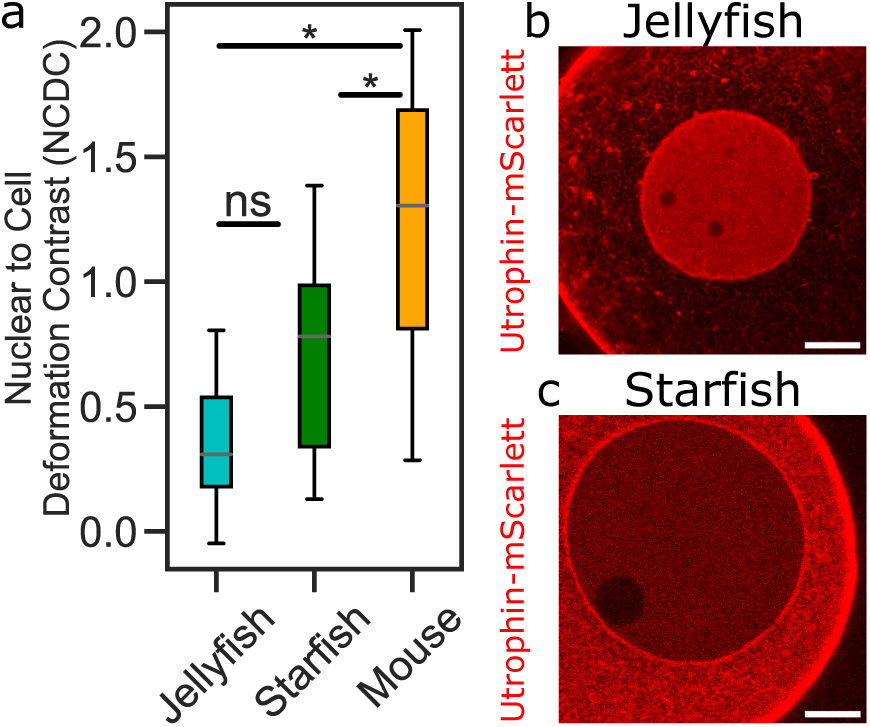
**a**: A comparison of the relative shape change of the nucleus *NCDC* between species. **b-c**: Live cell imaging nuclei with actin staining, in jellyfish and starfish (Utrophin-mScarletI3) oocytes. Scale bars: 20 *µ*m. (Jellyfish: *n* = 7; starfish: *n* = 15, mouse: *n* = 8. *: *p <* 0.05, ns: *p >* 0.05).

Motivated by these differences between two species, we wondered if the impressive shape mimicry is a special feature of starfish oocytes. To test this, we repeated the experiments with mouse oocytes. In these mammalian egg cells, we could observe that the shape mimicry is even stronger than in either starfish or jellyfish (Fig. 2a). A possible explanation is that in mice, the oocyte maturation and expulsion is fully shielded from forces that are originating from the outside of the organism, suggesting that, from a biological point of view, no mechanical protection mechanism are required. Still, during ovulation significant forces might be applied at follicle rupture,^33^ and the question if the effect of such deformations on the nucleus play a role for fertilization is intriguing.

### Dynamics of shape recovery after cellular compression

Although our study shows that the nucleus of jellyfish oocytes has a much lower *NCDC* than that of starfish, potentially due to nuclear actin, we wondered to which extend the observed deformations imprint a permanent deformation shape memory of the nuclei, or if these nuclear shapes are only transient. To test this we exploited the potential of the compression experiments to also provide access to the dynamical response of the cellular and nuclear shape. This is shown in Fig. 3a and Supplementary Movie 1, where *NCDC* is plotted as a function of time for both the nucleus and the whole starfish oocyte. The indentor is removed quickly, typically within the course of a single frame, so that the relaxation is purely due to the viscoelastic response of the cell and nucleus. The instantaneous relaxation characterizes the elastic response, while the viscous response takes place over the course of several seconds. Similar to other studies on the viscoelastic properties of cells,^37, 38^ we fitted the relaxation data with a power law:

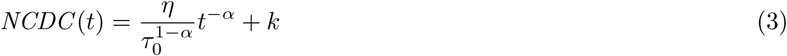

where the prefactor *η* is the magnitude of the viscous response, *k* an instantaneous elastic offset, *α* an exponent, and *τ*_0_ = 1 s is the reference timescale.

**Figure 3:**
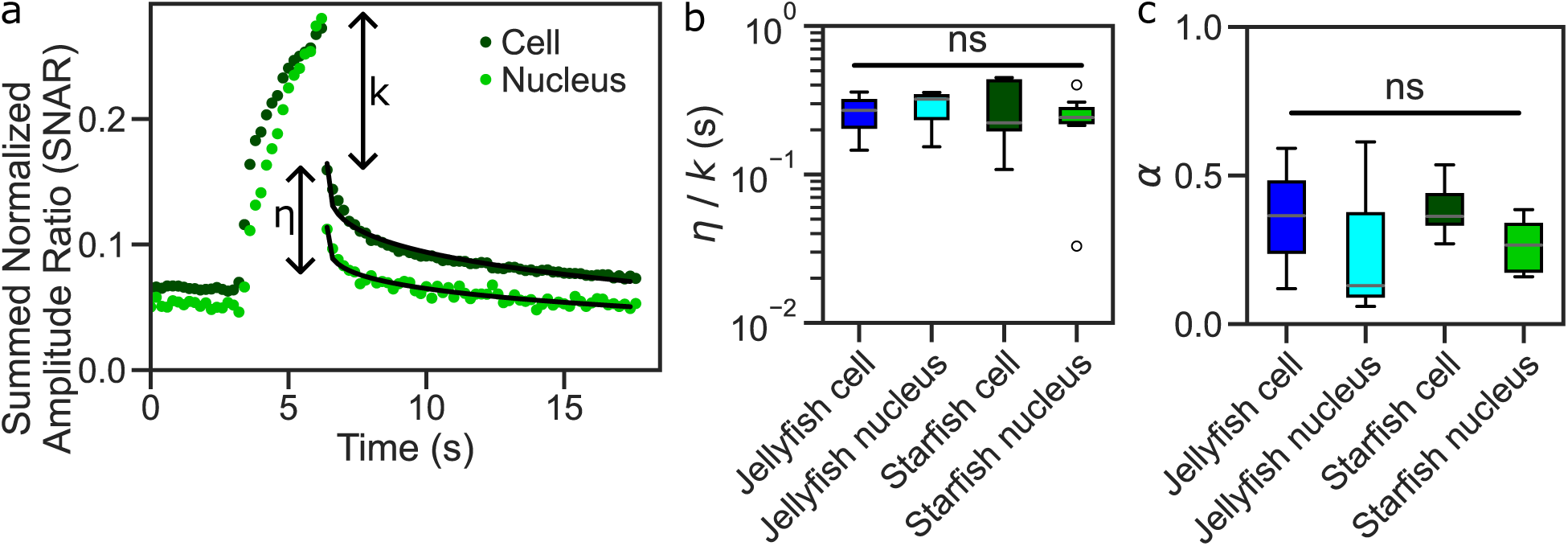
Time-dependent recovery of oocyte deformation. **a**: The 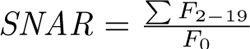 as a function of time for a starfish oocyte (red) and its nucleus (blue). After removal of indentation around *t* = 3 s, the cell and nucleus undergo viscoelastic relaxation, with an instantaneous, elastic response that is quantified by *k* and a viscous response *η* that was fitted by a power law (black line). **b**: Overview of the viscous timescale *η/k* for starfish and jellyfish oocytes. **c**: Overview of the exponent *α* for starfish and jellyfish oocytes. (starfish: *n* = 9, jellyfish: *n* = 3. No significant difference (*p >* 0.05) was found in panel **b** and **c**.)

In this view we can introduce a viscous timescale by the ration between *η* and *k*. This is shown in Fig. 3b for whole cells and nuclei. As the mouse oocytes (as well as some of the jellyfish oocytes) were sticking to the indentor, only data for starfish and jellyfish are shown. Additionally, Fig. 3c shows the power law exponent *α*. We observe two things from these graphs: first, the relaxation response is mostly elastic as *k* dominates (as indicated by *η/k <* 1), and secondly, the viscous timescale does not seem to differ between the whole oocyte and the nucleus. This is in line with a nucleoplasm that relaxes faster than the whole cell, and is only limited by the relaxation of its surroundings. Relaxation experiments of isolated nuclei could further demonstrate this point.^12, 18^

Interestingly, the deformability of the nuclei, being very similar to the deformability of the cell, and the undelayed time-dependent recovery both suggest that the viscoelastic properties of the nucleus are softer and less viscous than those of the whole cell. This is in stark contrast to earlier studies that reported high stiffness values of nuclei. A possible explanation for this discrepancy lies in one major difference with previous studies, namely that the size of the oocyte and its nucleus are an order of magnitude larger than that of previously studied cell types. Consequently, the ratio of the nuclear volume to its area differs by a similar amount, which in turn suggests that the mechanical role of the nucleoplasm with respect to the nuclear envelope is much larger. We thus hypothesize that the nucleoplasm is significantly softer and less viscous than the cytoplasm of oocytes, to account for the high deformability of the nucleus.

### Comparing cytoplasmic and intranuclear mechanics

To test the hypothesis that the oocyte nucleoplasm is softer than the surrounding cytoplasm, we directly measured the viscoelastic material properties of the starfish and jellyfish oocytes and nuclei. To this end, we injected passivated polystyrene particles with a diameter of 1 *µ*m into either the nucleus or the cytoplasm of jellyfish and starfish oocytes, and into the cytoplasm of mouse oocytes (Fig. 4a-b). These particles were used as probes to perform optical tweezers based microrheology that allows quantifying the frequency-dependent viscoelastic shear modulus *G*^∗^(*f*) (shown schematically in Fig. 4c). We find that over the entire measured frequency range from 1 Hz to 2.7 kHz both the storage and loss modulus *G*^′^(*f*) and *G*^′′^(*f*) are more than an order of magnitude higher for the cytoplasm than the nucleoplasm (Fig. 4d-e). Fig. 4f shows tan *δ* = *^G′′^* at 1 Hz, directly demonstrating that the nuclei are dominated by the viscous modulus, and hence considered a fluid (i.e. tan *δ >* 1) while in the viscoelastic cytoplasm the storage and loss modulus are similar (i.e. tan *δ ≈* 1).

**Figure 4:**
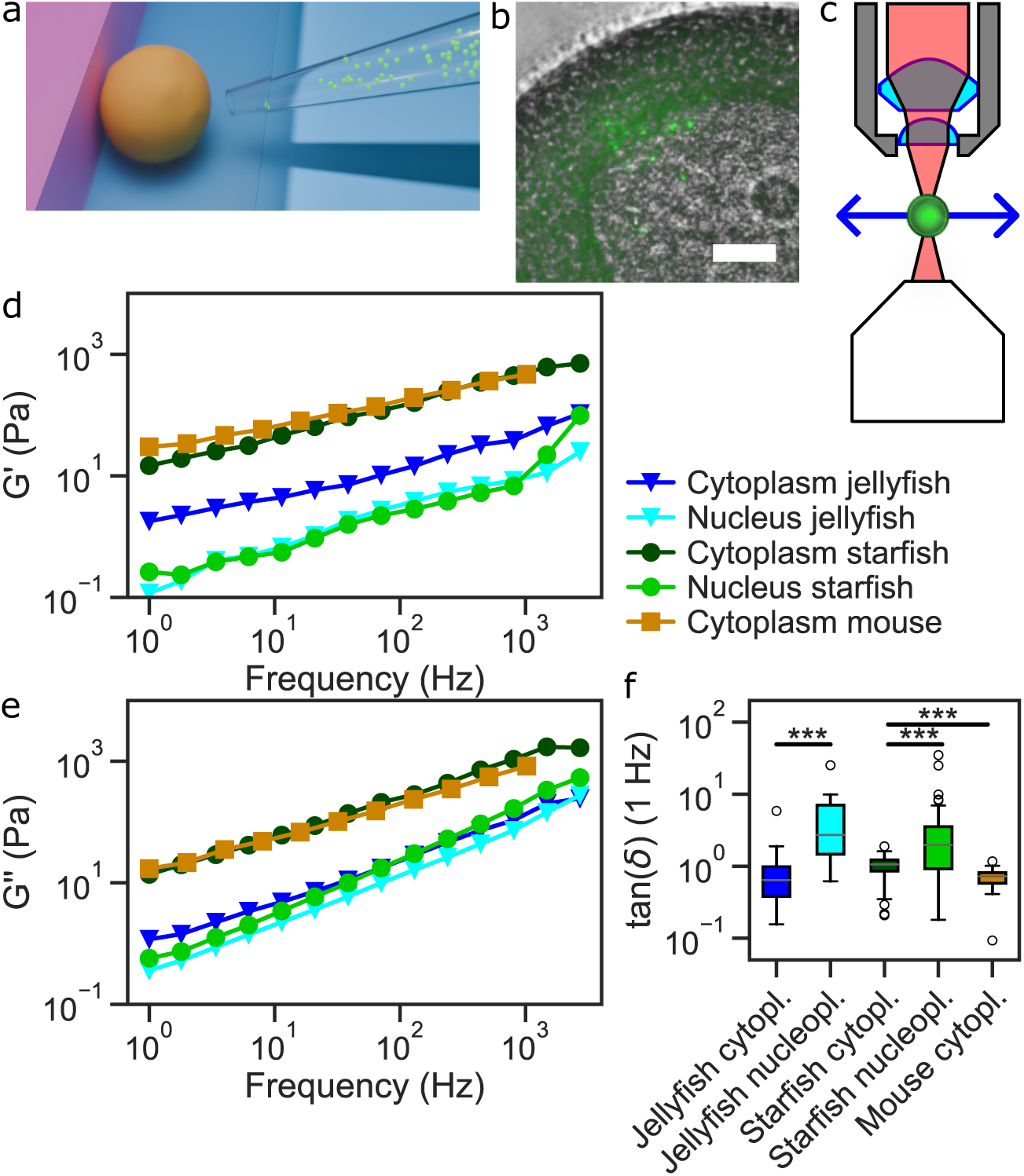
**a**: Schematic overview of oocyte injection with micron-sized polysterne beads using a micropipette and confinement of the oocyte between two glass cover slips, **b**: fluorescent image of an oocyte overlaid with a brightfield image, with 1 *µ*m injected beads clearly visible in the cytoplasm and nucleoplasm. Scale bar: 20 *µ*m. **c**: Schematic overview of the optical tweezers setup. **d-f**: Microrheology measurements of the **d** storage (*G*^′^) and **e** loss (*G*^′′^) moduli and **f** their ratio (tan *δ*) at *f* = 1 Hz, measured on oocytes of three different animal species. Curves are averages between *n* = 33 (jellyfish cytoplasm), *n* = 10 (jellyfish nucleus), *n* = 61 (starfish cytoplasm), *n* = 31 (starfish nucleus) and *n* = 11 (mouse cytoplasm) measurements. ***: *p <* 0.001.

To further quantify the differences between the mechanical properties of the nucleus and cytoplasm, we fitted the mechanical properties *G*^∗^(*f*) = *G*^′^(*f*) + *iG*^′′^(*f*) with a fractional viscoelastic model.^39–41^ The resulting model parameters quantify elasticity, viscosity and the level to which an elastic and fluid description is adequate (Supplementary Figure 1). Similar to the *G*^∗^(*f*) as show in Fig. 4d-e, the fractional viscoelastic model demonstrates significant differences between the properties of the cytoplasm and the nucleoplasm. Furthermore, we can identify for the fluid-like part of the shear modulus an exponent that is very close to 1, suggesting that the nucleoplasm behaves as a perfect Newtonian fluid surrounding an elastic component of chromatin and/or actin filaments.

Additionally, these measurements show that overall the jellyfish nucleoplasm is not only less viscous than the starfish nucleoplasm, the remaining elastic modulus is also smaller. While this does at first sight contradict the finding that jellyfish nuclei have a considerable concentration of actin filaments, it suggests that lower *NCDC* deformation value is either controlled by the nuclear envelope or by the lower cytoplasmic stiffness in the jellyfish. Microrheology measurements, however, are not probing the nuclear envelope, but only provide information on the nucleoplasm and the cytoplasm. Therefore we turned to Finite Element Method simulations to test whether the differences in deformability between starfish and jellyfish oocytes are better explained by the different cytoplasm mechanics or require a contribution of the nuclear envelope stiffness.

### Simulations of oocyte deformation

To test the hypothesis that the nuclear envelope in the jellyfish oocyte is much stiffer than in the starfish oocyte, we exploited the information about the mechanical properties of the cytoplasm and the nucleoplasm to simulate the system while varying the nuclear envelope mechanics until we obtain values that are in agreement with the measurements. Using a finite element approach (see Fig. 5a), we approximate the oocytes as a sphere that is positioned inside a sphere. For the cell boundary we use previous measurements: we assign a shell thickness of 1.4 *µ*m and 1 kPa Youngs modulus. For the nucleus we keep the thickness of the shell constant (0.3 *µ*m) and vary the stiffness. The mechanical properties of the cytoplasm and the nucleoplasm are adapted to represent the values we measured using active microrheology. As we here implement a static measure, we use the elastic modulus at 1 Hz. In details, that means for the starfish *E_cyto_^SF^* = 42.6 Pa, = 0.75 Pa and for the jellyfish *E_nucl_^JF^* = 5.2 Pa, *E_nucl_^JF^* = 0.35 Pa. The simulation then enforces the indentation as applied in the experiment. Hence, variations in the stiffness and thickness of the cell membrane affect the deformation of the nucleus very little, as the applied deformation is strain-controlled.

**Figure 5:**
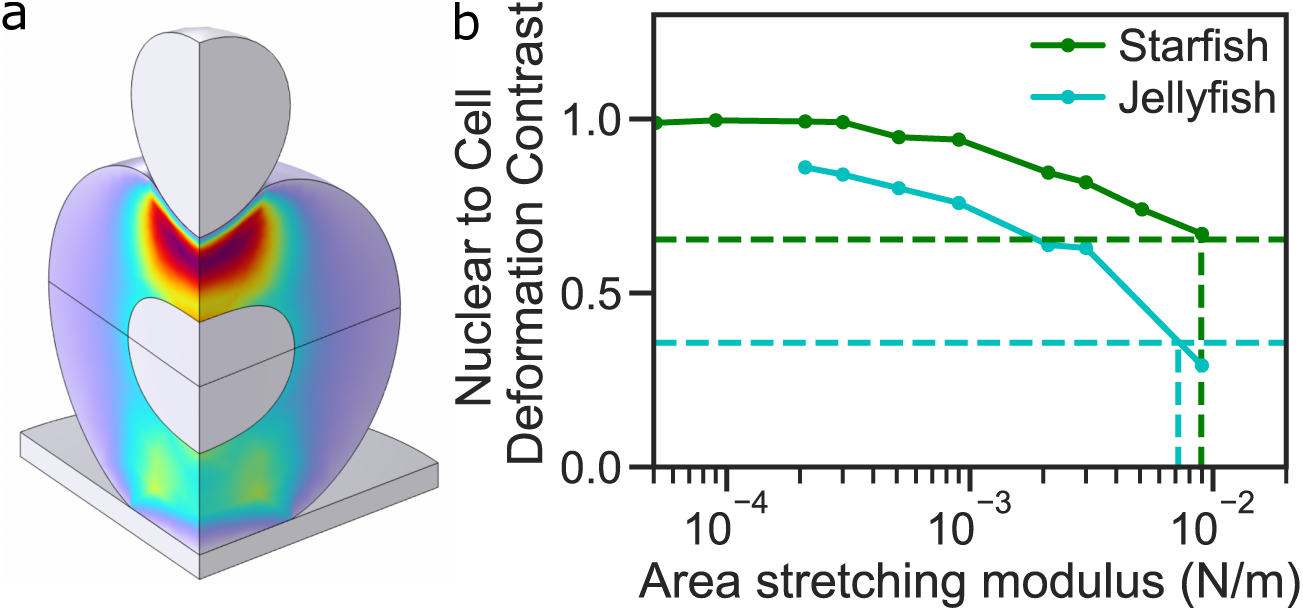
Simulations of oocyte deformation **a**: Snapshot of the simulated oocyte and nucleus during indentation, and **b**: the Nuclear to Cell Deformation Contrast (*NCDC*) as a function of the area stretching modulus (the product of the stiffness and thickness of the nuclear envelope with the lamin network). Dashed lines indicate the experimentally observed values for starfish (green) and jellyfish (cyan) oocytes.

By varying the stiffness of the nuclear envelope we obtain the *SNAR* as defined in Eq. 1 as a function of the nuclear envelope area streching modulus, which is calculated *K_A_* = *E · d*, with the Youngs modulus *E* and the thickness of the shell *d* = 0.3 *µ*m. In this system the stretching modulus becomes independent of the thickness, which makes it a reasonable mechanical number to extract. Fig. 5b shows resulting *NCDC* as a function of the used area stretching modulus. In this plot the green dashed line indicates the *NCDC* that we find experimentally for starfish oocytes and the cyan line the *NCDC* for jellyfish oocytes. Indeed, we find here a reasonable value for the area stretching modulus of *K_A_^SF^ ≈* 9 *·* 10^−3^*N/m*, and *K_A_^JF^ ≈* 7 *·* 10^−3^*N/m* respectively. Such values are in a similar order of magnitude as reported for the cell cortex of somatic cells.^42^ Since the area stretching moduli of the different nuclei are quite similar, this result suggests that the differences in *NCDC* is due to the different cytosolic mechanical properties, and not related to changes in the nucleus or in the nuclear envelope.

## Discussion

### Oocyte nuclear mechanics differ from non-germ cells

The surprising result that oocyte nuclei follow shape changes applied to the overall cell is a special property of oocytes. The nucleoplasm behaves as a liquid, and although the cytoplasm is quite soft, its mechanical properties are responsible for the propagation of external strains to the nucleus, which in turn deforms in a way that mimics the global shape changes. Different species show this effect to different degree, where cells with a softer cytoplasm show less nuclear deformation.

A reasonable mechanistic explanation lies in their sheer size. Oocyte cell and nuceli volume are almost an order of magnitude larger than their somatic counterparts, and the ratio of the nuclear volume to its area differs by a similar amount. Consequently, this increases the mechanical role of the nucleoplasm with respect to the nuclear envelope. As the nuclear envelope is mechanically shielding the nucleoplasm, a reduction in its relative contribution leads to more deformable nuclei in oocytes. Additionally, compared to somatic cells, the amount of chromatin in the oocytes remains the same while the nuclear volume is about 1000 times larger in the oocytes. Therefore, the chromatin to which the mechanical properties are commonly attributed is diluted by *≈* 10^3^, thus making the nucleoplasm softer.

### Comparing internal versus external mechanics

By looking in more detail at the viscoelastic nature of the nucleoplasm and the cytoplasm versus the whole nucleus and oocyte, we find an apparent discrepancy. Step strain and subsequent relaxation experiments show that the whole oocytes and nuclei respond mainly elastic to external deformations. In contrast, the active microrheology data does not confirm this, as the intranuclear environment is predominantly viscous, and the cytoplasm is almost perfectly viscoelastic. Additionally, the resistance to deformations as found in jellyfish cannot be explained by the nucleoplasm, since it is much softer than the surrounding cytoplasm.

To understand this discrepancy between the microrheology data and the different deformations of the nucleus in the tested species, we need to consider the nucleus as a multiphasic material, consisting of a mainly viscous nucleoplasm and a mainly elastic nuclear envelope.^10^ The origin of the elasticity is the network of lamin fibers that line the inside of the nuclear envelope.^43^ All animal species used in this study express lamin in their oocytes: most invertebrate oocytes only have one type of lamin,^44^ whereas mammalian oocytes express both lamin A, B and C2.^45^

Hence, intranuclear mechanics play only a minor role when the nucleus is probed from outside as the nuclear envelope and the lamin network essentially shield the nucleoplasm from deformations. This view of a biphasic nucleus consisting of an elastic shell and a soft interior is further supported by the simulations of the indentation that hint at a strongly elastic response of the nuclear envelope and the underlying lamin network.

A similar apparent contradiction between internal and external mechanical measurements exists in measurements of cellular stiffness. Measurements on cellular stiffness from the outside of a cell report values ranging from 1 to 100 kPa,^46^ while intracellular measurements typically show values between 10 and 100 Pa (Fig. 4 and Refs.^39, 47^). This discrepancy is due to the actin cortex that is found directly under the cellular plasma membrane. Hence, external measurements predominantly probe the stiff and elastic actin cortex, while intracellular measurements directly probe the viscoelastic cytoplasm.

The view that the nucleoplasm is relatively soft is supported by the observation that the recovery of the nuclear deformation instantaneously follows cellular recovery, as seen in Fig. 3 and Refs.^6, 16^ The viscous component of the mechanical response dictates the speed by which the shapes of the cell and the nucleus recover after deformation. As the nuclear shape recovers simultaneously with the cell shape upon strain removal, this indicates that the viscosity of the nucleus is equal or lower than the viscosity of the cytoplasm, in direct agreement with our microrheology results (Fig. 4b). This is further supported by Refs.,^48, 49^ where it was found that the refractive index and mass density of the nucleus is lower than the cytoplasm in certain human cells, and Ref.,^50^ which shows that the nucleus of *Xenupus* egg extract has a lower mass density than the surrounding cytoplasm. Although refractive index and mass density do not directly correlate to mechanical properties, it fits a picture of fewer dissolved proteins and small organelles, and hence a lower viscosity.

But why does the jellyfish nucleus deforms less upon whole cell deformation? As the simulations show, the contribution of the nuclear envelope is comparable between jellyfish and starfish. However, since the cytoplasm in the jellyfish cells has a much smaller elastic modulus compared to starfish and mice, it also transmits less of the externally applied strain, thus explaining that the jellyfish nucleus follows much less the global cell shape as in the other two species. Here, the reduced actin concentration in the cytoplasm seems to be the main reason for the lower elasticity. However, it remains a surprise that the nuclear stiffness stays the same when comparing starfish and jellyfish despite the drastic increase of actin filaments in the starfish oocytes.

### Implications for cell signalling

All in all, the mechanical properties of oocytes and their nuclei are profoundly different from those of most cell types in starfish, jellyfish and mouse. The nuclei are more prone to deformations, which, given the importance of mechanical signalling, can have profound consequences on a biological level. Although cells adapt to local deformation, the time scale for remodeling of the fibrillar adhesions that anchor the vimentin cytoskeleton is on the order of an hour,^51^ suggesting that active remodeling does not occur on the time scale of our applied indentation.

## Conclusion

In this study, we found little resistance to deformation of the nucleus when deforming the full cell. Albeit the effect of nuclear shape mimicry is species dependent, the effect can be observed in mouse, starfish and jellyfish. We explain this by a softer and less viscous nucleoplasm compared to the cytoplasm. This hypothesis was confirmed by performing microrheology on particles injected in both the cytoplasm and the nucleoplasm. We found that the elastic and viscous properties of the nucleus were at least an order of magnitude softer than in the cytoplasm. This means that in oocytes, the nuclei are not stiff, but soft. This finding can be understood by the different chromatin density in oocytes, which is diluted by about a thousand times, as well as a change in the high volume-to-surface ratio due to the large cell size.

## Materials and Methods

### Oocyte harvesting

Starfish: (*Patiria miniata*) ovaries were obtained through a biopsy on the dorsal side of an arm using a surgical biopsy punch (size 3, kai medical). The extracted pieces of ovary were placed for 20 to 30 minutes in ice-cold calcium-free filtered sea water (FSW, pH 8) containing 50 mM L-Phenylalanine (Sigma) to prevent spontaneous oocyte maturation. Subsequently, the ovaries are transferred to FSW containing 100 *µ*M Acetylcholine (Sigma). This induces contraction of the ovaries, leading to the expulsion of oocytes from the ovary. The oocytes are then maintained in Petri dishes with FSW at 14°C for up to 3 days.

Jellyfish: Each fully grown female *Clytia hemisphaerica* jellyfish contains four ovaries with oocytes at various developmental stages. Ovaries are surgically isolated from the jellyfish the day prior oocyte isolation. Isolated ovaries are then kept in FSW at 18°C, and are subjected to an artificial light cycle lasting approximately 16-20 hours to facilitate the development of oocytes to their final stage. On the next day, fully grown oocytes are isolated by carefully cutting open the ovary and extracting the oocytes using a glass needle. The isolated fully-grown oocytes are placed in a Petri dish containing FSW to be used on the same day.

Mouse: Maintenance and handling of all mice were carried out in the MPI-NAT animal facility according to international animal welfare rules (Federation for Laboratory Animal Science Associations guidelines and recommendations). Requirements of formal control of the German national authorities and funding organizations were satisfied, and the study received approval by the Niedersächsisches Landesamt für Verbraucherschutz und Lebensmittelsicherheit (LAVES). All mice were maintained in a specific pathogen-free environment according to The Federation of European Laboratory Animal Science Associations guidelines and recommendations. Mice are housed in individually ventilated cages at 21°C and 55% relative humidity with open cage changing. Animals are provided with commercial soy-free breeding chow and water ad libitum.

### Oocyte preparation and injection

For compression experiments, oocytes were put on a 22×22 mm cover slip glued on top of a 24×60 mm microscope slide. A droplet of sea water (starfish and jellyfish oocytes) or medium (mouse oocytes) prevented evaporation. For microrheology experiments, oocytes are mounted in between two glass coverslips separated by a spacer of double-sided tape. This configuration ensures that the oocytes are immobile during injection and microrheology measurements.

For fluorescence labeling in compression experiments, Dextran labeled with Tetramethylrhodamine isothiocyanate (TAMRA/TRIC/DEXTRAN, 155 kDa, Sigma T1287) was dissolved in ddH2O to 25 mg/ml. From this stock, a 1:10 dilution in ddH2O was utilized for microinjections of starfish oocytes. Jellyfish oocytes naturally express GFP, hence no fluorescence labeling is required.

For micorheology experiments in starfish and jellyfish oocytes, 1 *µ*m fluorescently labeled polystyrene beads (Micromod, Rostock, Germany) were injected. The beads were coated with PEG to avoid bead aggregation and adhesion to the inner walls of the micropipette during injection, as well as to ensure that there are no specific interactions of the beads with nuclear or cytoplasmic components. Briefly, beads were sonicated and washed in a borate buffer, before adding MS(PEG)12 (Fisher Scientific GmbH, Schwerte, Germany) and let to incubate for 2 hours at 4°C, after which the beads were washed again.

Injection was performed with home-made glass capillaries (Cat#1-000-0500, Drummond scientific company, Broomall, USA). Prior to injection, beads were sonicated to avoid clustering and diluted 1/10 to 1/20 in PBS. The bead suspension was loaded into the back of the micropipette using a flexible needle. The tip was broken by gently pushing the needle against the side of a glass coverslip.

Mouse oocytes for bead injection experiments were isolated from ovaries of 8-12-week-old female FVB/N mice. Fully grown GV (germinal vesicle) oocytes (*≈* 75 *µ*m in diameter) with a central nucleus were selected for experiments and cultured in homemade M2 medium supplemented with 250 *µ*M dibytryl cyclic AMP (dbcAMP) (Sigma Aldrich, #D0627) covered with paraffin oil (NidaCon #NO-400K).

For bead injection in mouse oocytes, 1 *µ*m fluorescent beads were prepared as described above. Oocytes were injected on a FemtoJet injection system (Eppendorf) with Piezo (Eppendorf) mounted on an Axio Vert.A1 inverted microscope (Zeiss) using a DIC 40X air objective (Zeiss). Oocytes were transferred to a 35 mm, low uncoated Ibidi imaging dish (Ibidi, #17266143) to accommodate the angled injection needle and holding pipette. Injection needles were prepared from glass capillaries with a diameter of 1 mm. The needle tips were bent to an angle of 30°. Holding pipettes (Eppendorf VacuTip I, #EP5195000036) were pre-filled with medium and connected to a CellTram Air pump (Eppendorf). Injection needles were backloaded with the prepared bead solution. The injection needles were connected to a FemtoJet 4i system (Eppendorf). Oocytes were loaded into a large drop of M2 medium supplemented with dpcAMP to maintain the GV arrested stage. Beads were loaded into an injection needle with a bent tip, and oocytes were immobilized with the holding pipette. The FemtoJet was adjusted to ensure a steady flow of beads. Oocytes were injected by piercing the membrane and injecting beads with a short pulse. On average, 2-10 beads were injected. Oocytes were maintained at 37°C throughout the experiment.

For microrheology experiments, oocytes were transferred to a coverslip and M2 medium was carefully replaced with low-melting agarose dissolved in warm M2 medium. Care was taken that oocytes did not move, maintaining the position on the coverslip. The temperature of the low-melding agarose was reduced to 37°C just prior to applying to oocytes to avoid heat shock. The oocytes in agarose were covered with an additional coverslip and sufficient distance between the coverslips was ensured by separating the two coverslips with two layers of double-sided tape.

### Oocyte compression

An indentor with a blunt spherical cap was created by breaking off the tip of a micropipette and holding the remainder over a glowing metal wire to melt the opening and form a sphere with a radius of *≈* 40 *µ*m. This blunt tip was used for controlled cell deformation by pushing a free-floating oocytes against the side of a glass coverslip. Fluorescence (jellyfish and starfish) or brightfield (mouse) images were recorded at a rate of 5 frames per second with a Hamamatsu Fusion camera with a 10x air objective on an inverted Nikon Eclipse Ti2 microscope.

The contour of the starfish and jellyfish oocyte and nucleus was automatically detected with a thresholding method in a custom Python script. The contour of the mouse oocyte and nucleus was obtained manually in FIJI. To quantitatively analyse the shape of the nucleus, a Fourier transformation of the angle-dependent radius was taken, and the ratio was taken of the sum of second to the twentieth components to the zeroth component (corresponding to the radius of a perfect circle), similar to 2D^34, 36, 52^ and 3D^53^ approaches described previously.

### Labelling F-actin

Starfish oocytes were injected with Utrophin-mScarletI3 at a concentration of 484 *µ*M. Jellyfish oocytes were injected with Utrophin-mScarletI3 at a concentration of 125 *µ*M.

### Optical tweezers setup

Optical tweezers based microrheology was performed on a home-built setup as described elsewhere.^31, 47^ Briefly, the setup uses two infrared lasers to apply a force on a probe particle and detect its position, respectively. An 808 nm, 250 mW laser (Lumics GmbH, Berlin, Germany), is used to trap particles, as 808 nm infrared light is only minimally absorbed in aqueous environments compared to the more commonly used 1064 nm lasers, reducing local heating and photodamage. The trapping laser can be steered by a mirror glued onto a piezo tilting platform (S-331, Physik Instrumente (PI) GmbH & Co. KG, Karlsruhe, Germany). A second, 976 nm, 500 mW laser (Thorlabs, New Jersey, USA) is used at low power (typically 0.3 mW) for position detection. Both lasers are focused by a high NA objective (CFI Plan Apochromat VC 60XC WI NA 1.2, Nikon) to a diffraction-limited spot in the sample plane.

On the opposing side of the sample, the IR light is collected by a 1.4 NA oil immersion condenser and imaged onto a Position Sensitive Detector (976 nm laser: order nr. 5000011, First Sensor, Berlin, Germany) and onto a Quadrant Photo Diode (808 nm laser: QPD, PDQ80A, Thorlabs, New Jersey, USA) for the trapping and detector signal, respectively.

Sample positioning is achieved by a piezo stage with a 200 *µ*m travel range combined with a stepper motor (Mad City Labs, Kloten, Switzerland). The microscope allows both brightfield and epifluorescence imaging.

### Performing microrheology experiments

Active microrheology was performed by applying an oscillatory force to the particle using an optical tweezers as described previously.^31, 37, 39^ A frequency sweep was performed with driving frequencies *f_d_* evenly spaced on a logarithmic scale between 1 Hz and 2.7 kHz. By taking the Fourier transform of the force *F*^∼^(*f_d_*) and position signal *x*∼(*f_d_*), the response function *χ*^∗^ is calculated:

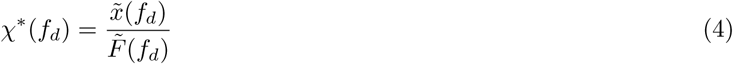

The complex shear modulus *G*^∗^(*f*) = *G*^′^(*f*) + *iG*^′′^(*f*) is then obtained by the relation:

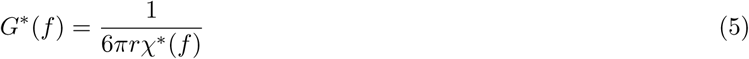

with *r* the radius of the (spherical) probe particle.

Averaging of *G*^′^, *G*^′′^ is done in log-space: for every driving frequency, the logarithm of all data points is taken first, then an average is taken per frequency and finally the exponent of the average is calculated, to return to linear space. By doing this, large outlier values do not disproportionally affect the average.

### Simulations of oocyte deformation

COMSOL Multiphysics 6.2 was used to simulate the mechanical response of starfish oocytes to compression (Supplementary File 1). The oocyte was represented by two concentric spheres. The inner sphere’s volume and surface represent the nucleoplasm and nuclear membrane, while the volume of the outer surface, excluding the nucleus, along with its surface, mimics the cytoplasm and cell cortex, respectively. The radii of nucleus and cell were 37.5 *µ*m and 85 *µ*m respectively, as determined from starfish oocyte images. Finally, the indentor was represented by a sphere of radius 40 *µ*m.

To model both the nucleoplasm and cytoplasm, we used a Neo-Hookean hyperelastic material for both the nucleoplasm and cytoplasm, since the indentation was comparable to the size of the nucleus. Additionally, we incorporated the viscoelasticity using a Kelvin-Voigt model. In this model, the Young’s modulus (*E*) and the relaxation time are required. Given that the experiments operated on a time scales of seconds, we used the value of storage modulus (*G*^′^) at 1 Hz as an estimate of the shear modulus (*G*). Then, assuming a value of 0.45 for Poisson’s ratio (*ν*), we could estimate the Young’s modulus (*E*) using Eq. 6. Furthermore, we utilized loss modulus (*G*^′′^) at 1 Hz to calculate the viscosity using Eq. 7 and subsequently determined the relaxation time using Eq. 8.

Due to the low thickness of the nuclear membrane and cell cortex in comparison to the cell size, we opted for the membrane interface for both membranes. For modelling the membranes, we chose linear elastic membranes, with the Young’s modulus of both membranes as the adjustable parameters in the simulation.

To model the indentation, a solid spherical indentor was stepwise moved downwards by 40 *µ*m. Given that the relaxation time of the materials is one order of magnitude shorter than the experimental time scale, the cell can quickly respond to the applied deformations. Hence, we used a stationary solver.

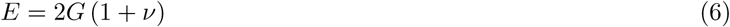

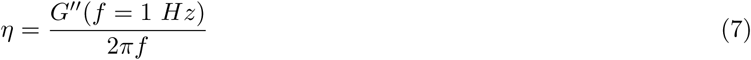

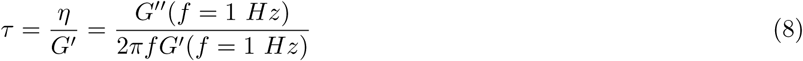

### Statistical analysis and data representation

Box plots extend from the first to the third quartile values of the data, with a line at the median. The whiskers extend from the box to the furthest data point in the 1.5x interquartile range. Flier points are those past the end of the whiskers.

Statistical analyses were performed using a two-sided Mann-Whitney *U* test.

## Data availability

All experimental data is available from the authors upon request.

## Code availability

All code will be made available from public repositories upon publication.

## Acknowledgements

BV, TM and TB have received funding from the European Research Council (ERC) under the European Union’s Horizon 2020 research and innovation programme (PolarizeMe, Grant agreement No. 771201) and by the Deutsche Forschungsgemeinschaft (DFG) under Germany’s Excellence Strategy (EXC 2067/1-390729940). Furthermore, TB, PL and MS were funded by the DFG – Project-ID 449750155 – RTG 2756. Research in PL’s laboratory is additionally supported by the Deutsche Forschungsgemeinschaft-Agence Nationale de la Recherche (DFG-ANR) project DOLLI (grant Nr. LE 2926/3-1). I.M.A.J. was supported by a Boehringer Ingelheim Fonds PhD fellowship. I.M.A.J. has been a doctoral student of the Ph.D. program “Molecular Biology” – International Max Planck Research School and the Göttingen Graduate School for Neurosciences, Biophysics, and Molecular Biosciences (GGNB) (DFG grant GSC 226) at the Georg August University Göttingen. We thank Jasmin Jakobi for help with the starfish handling, and Kerstin von Roden with coating of the polystyrene particles.

## Competing interest

The authors declare no competing interest.

## Supplementary Information

### Oocyte indentation

Supplementary Movie 1: indentation of a starfish oocyte.

### Fitting microrheology data with a fractional viscoelastic model

To quantify the differences between the mechanical properties of the nucleus and cytoplasm, we fitted *G*^∗^(*f*) = *G*^′^(*f*) + *iG*^′′^(*f*) with a fractional viscoelastic model^39, 41^

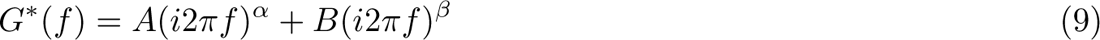

Confirming the frequency-dependent plots, we find strong significant differences in the fitting parameters of the cytoplasm and the nucleus, see Supplementary Fig. 1. The prefactors *A* and *B* are more than an order of magnitude larger for the cytoplasm than for the nucleus. The value of the power law exponent *α* is close to 0.5 for the cytoplasm, reminiscent of an elastic environment, while for the nucleus the exponent *β* is very close to 1, resembling an almost completely liquid environment.

### Finite Element simulation

Supplementary File 1: COMSOL-file of the oocyte under deformation.

---

**Supplementary Figure 1:**
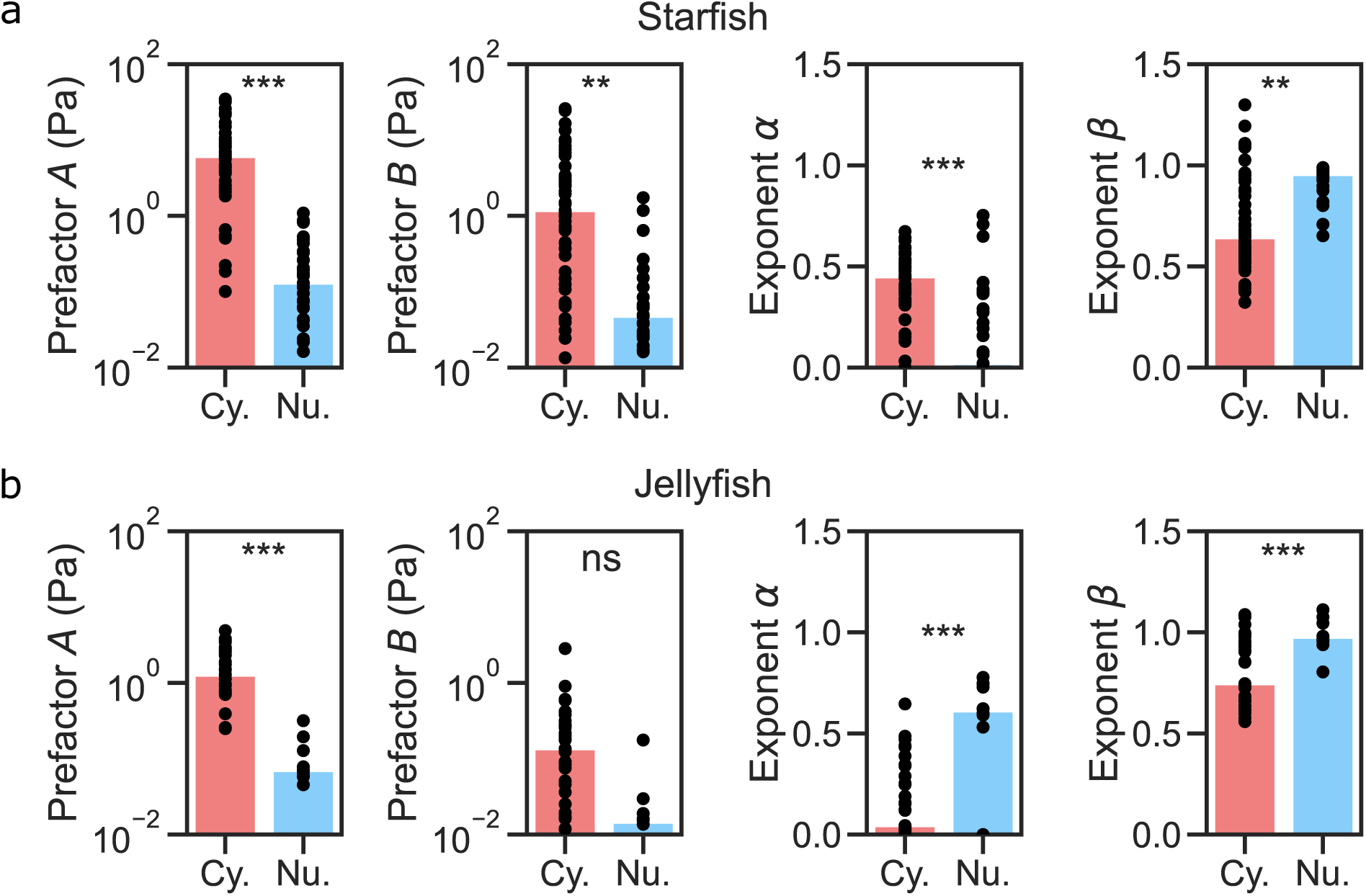
Fitting parameters of *G*^′^ and *G*^′′^ with a fractional viscoelastic model. **Top**: starfish oocytes, and **bottom**: jellyfish oocytes. The bar indicates the median value of the fit parameter. (***: *p <* 0.001; **: *p <* 0.01, ns: *p >* 0.05)

## Notes

### Competing Interest Statement

The authors have declared no competing interest.

